# Single Nucleotide Polymorphisms in the Bovine TLR2 Extracellular Domain Contribute to Breed and Species-Specific Innate Immune Functionality

**DOI:** 10.1101/2021.08.23.457326

**Authors:** Marie-Christine Bartens, Amanda J. Gibson, Graham Etherington, Federica di Palma, Angela Holder, Dirk Werling, Sam Willcocks

## Abstract

Recent evidence suggests that several cattle breeds may be more resistant to infection with the zoonotic pathogen *Mycobacterium bovis than others*. Our data presented here suggests that the response to mycobacterial antigens varies in macrophages generated from Brown Swiss (BS) and Holstein Frisian (HF) cattle, two breeds belonging to the *Bos taurus* family. Whole genome sequencing of the Brown Swiss genome identified several potential candidate genes, in particular Toll-like Receptor-2 (TLR2) a pattern recognition receptor (PRR) that has previously been described to be involved in mycobacterial recognition. Further investigation revealed single nucleotide polymorphisms (SNP) in TLR2 that were identified between DNA isolated from cells of BS and HF cows. Interestingly, one specific SNP, H326Q, showed a different genotype frequency in two cattle subspecies, *Bos taurus* and *Bos indicus*. Cloning of the TLR2 gene and subsequent gene-reporter and chemokine assays revealed that this SNP, present in BS and *Bos indicus* breeds, resulted in a significantly higher response to mycobacterial antigens as well as tri-acylated lipopeptide ligands in general. Comparing wild-type and H326Q containing TLR2 responses, wild-type bovine TLR2 response showed clear, diminished mycobacterial antigen responses compared to human TLR2, however bovine TLR2 responses containing H326Q were found to be partially recovered compared to human TLR2. The creation of human:bovine TLR2 chimeras increased the response to mycobacterial antigens compared to the full-length bovine TLR2, but significantly reduced the response compared to the full-length human TLR2. Thus, our data, not only present evidence that TLR2 is a major PRR in the mammalian species-specific response to mycobacterial antigens, but furthermore, that there are clear differences between the response seen in different cattle breeds, which may contribute to their enhanced or reduced susceptibility to mycobacterial infection.

## Introduction

Early recognition of pathogens and activation of the innate immune response is critical in determining the outcome of infection. Pathogens are recognized by evolutionary conserved, germline-encoded and cell-surface expressed pattern recognition receptors (PRR)[1]. One of the major groups of PRRs are the toll-like receptors (TLR), ten of which have been discovered common to both bovine and human immune systems [2–5] and are expressed on myeloid antigen presenting cells such as macrophages (MØ). Each of these TLR has evolved to recognise specific pathogen associated molecular patterns (PAMP) – and docking of these ligands to the extracellular domain (ECD) of the TLR triggers an intracellular signalling cascade [6]. This signalling is mediated by the recruitment of adaptor proteins to the intracellular Toll/interleukin-1 receptor homology (TIR) domain of the TLR [7,8]. While the amino acid sequence of the ECD varies considerably between species [9], the TIR domain is highly conserved [7,9].

Although TLR4, and possibly TLR9 and TLR8 are capable of sensing mycobacteria [10–12], recognition of mycobacterial antigens by TLR2 has been shown to be a key factor in determining disease progression in tuberculosis [12–14] [15]. [14,16]. [17] [18]. TLR2 recognizes a range of mycobacterial cell-wall antigens including the 19-kDA mycobacterial lipoprotein and glycolipids such as lipoarabinomannan (LAM), lipomannan (LM), the 38kDA antigen, LprG lipoprotein and phosphatidylinositol mannoside (PIM) [11,19,20]. Additionally, secreted mycobacterial proteins have been shown to act as TLR2 ligands. For instance, the well-studied virulence factor ESAT-6 has been shown to inhibit NF-κB activation in a TLR2-mediated manner [21]. Similarly, recombinant antigen TB10.4 from *M. bovis*, encoded by a subfamily of ESAT-6 and also present in *M. tuberculosis* and *M. bovis* BCG [22], has been shown to induce pro-inflammatory cytokine production via a TLR2 mediated NF-κB pathway [23].

Engagement of TLR2 expressed on MØ by mycobacteria can result in complex outcomes that can either promote or inhibit a protective inflammatory response. For example, TLR2 activation can result in the upregulation of pro-inflammatory cytokines within a variety of immune cells, at the forefront MØ. Upregulated cytokines include IL-1β, TNFα and IL-6; as well as reactive oxygen species (ROS) production through activation of NF-κB and MAPK pathways [24–26]. Furthermore, TLR2 signalling regulates the expression of iNOS, which interacts with NADPH oxidase and ROS to create reactive nitrogen intermediates [27,28] and can induce autophagy [29] - all essential mechanisms in cellular host defence to mycobacterial infection [30,31]. On the other hand, engagement of TLR2 by the secreted mycobacterial antigens, heat-shock protein (hsp) 60 and proline-proline-glutamic acid (PPE)18 initiates an anti-inflammatory response via IL-10 production [32,33]. In addition, ESAT-6 has been shown to inhibit IL-12 production upon binding to TLR2 in RAW cells [21]. Further immunomodulatory mechanisms of some mycobacterial species include the inhibition of MHCII receptor expression and therefore presentation of mycobacterial lipoprotein antigens [34,35]. Thus, mycobacteria can utilise TLR2 to manipulate the immune outcome and promote survival [15,36][13,14,19,21,29,36].

Given the importance of TLR2, it is perhaps not surprising that polymorphisms within the TLR2 gene have been shown to affect the immune response to mycobacterial infection in both humans and cattle [18,29,32,33,37,38], [12,14]. In humans, the R753Q and R677W mutations have been shown to influence susceptibility to tuberculosis in case-control studies [37,39]. Furthermore, the R753Q SNP was also shown by Pattabiraman *et al* [40] to result in impaired NF-κB signalling and decreased cytokine responses to *M. smegmatis* in a murine and Human Embryonic Kidney (HEK) 293 cell assay. Similar as in the human system, several SNPs have also been identified within the bovine TLR2 orthologue. Some of these have been indicated to contribute to disease susceptibility in cases of paratuberculosis [41], clinical mastitis [42,43], and bovine tuberculosis [44,45][46]. SNPs occurring between cattle breeds may reflect natural variation arising from host-pathogen co-evolution in different geographical environments, but may also be influenced by intensive selective breeding that favours meat or milk production [9,47]. Characterisation of SNPs arising in TLR2 among commercially important breeds is therefore a crucial contribution towards understanding immune fitness variation among herds and potentially, disease sensitivity. While some SNPs within the bovine *tlr2* gene have previously been identified, studies of their functional relevance are often lacking. In the present work, we compare the *tlr2* sequences of two breeds of global importance to the farming/dairy industry, Brown Swiss (BS) and Holstein Friesian (HF) and determine the genotypic frequency of a H326Q SNP that occurs in the ligand-binding region of the ECD. We report on the species- and breed- specific phenotypic consequence of this mutation in response to canonical TLR2 ligands and in the context of challenge with *M. tuberculosis* and *M. bovis*. We present evidence that TLR2 is a major PRR in the mammalian species-specific response to mycobacterial antigens, that there are clear differences in species-specific responses that may influence disease susceptibility and host tropism of *Mycobacterium tuberculosis* complex members.

## Methods

### Mycobacterial Strains and Culture

The following strains were used for all assays as indicated: *M. bovis* strain AF2122/97 and *M. bovis* BCG Tokyo; *M. tuberculosis* H37Rv. Strains were cultured in BD Middlebrook 7H9 media (BD Difco™, USA), supplemented with OADC and 0.05 % Tween 80 (Sigma-Aldrich, UK). For *M. bovis* culture only, growth media was additionally supplemented with sodium pyruvate; for *M. tuberculosis* culture only, media was additionally supplemented with glycerol (Sigma-Aldrich, UK).

### Brown Swiss genome assembly and QC

Four lanes of 10x Genomics data were generated using an Illumina HiSeq. This provided ~1200 million reads with a 150bp read length. The data was assembled *de novo* using the 10x Genomics assembly tool Supernova v2.0.1, using the option to output a phased genome, which generates two homologous assemblies where each scaffold in the assembly has one of the paternal or maternal haplotypes [48]. Using one of the haplotypes, we used RepeatMasker, using ‘bovidae’ as the repeat library to soft-mask repeat regions in the assembly [49]. We tested the assembly for completeness using several quality control metrics. First, BUSCO (v5.0.0) was used to identify the number of mammalian single-copy orthologs present in the genome (using the database mammalia_odb10) [50]. Using the Kmer Analysis Toolkit, we examined the content and distribution of kmers in the assembly to that of the reads using the tool ‘kat comp’, ignoring the first 16 bases of each R1 read in order to omit 10x barcodes from the analyses. We then visualised the distribution of kmers using the tool ‘kat plot’ [51].

### Brown Swiss genome annotation and gene expansion

LiftOff tool was used to lift over the *Bos taurus* genome annotation (ARS-UCD1.2.103) to that of the Brown Swiss, noting any genes that were unmapped in Brown Swiss [52]. We then identified genes in the Brown Swiss genome annotation that were present in multiple copies (compared to *Bos taurus* ARS-UCD1.2.103) and downloaded the functional information for these genes using ENSEMBL BioMart database [53]. For genes annotated as being duplicated three or more times and where a gene name or description was missing, we downloaded the coding sequence for that gene and used megablast to search the NCBI nr database and noted the highest scoring hit (with 100% query coverage and an E-value equal to 0), that contained functional information about the gene (i.e., disregarding hits for clones, BACs, isolates, etc.).

### Study Population for Candidate Gene TLR2 analysis

Complementary DNA (cDNA) from TLR2 mRNA sequences isolated from PBMCs from six clinically healthy pedigree BS and four HF were investigated for sequence comparison. All cows were female and either housed, under Home Office License PPL7009059. Age and lactation cycle of sampled cows were recorded. After identification and selection of candidate SNP H326Q within the coding sequence (CDS) of TLR2 in the BS samples, further cDNA was generated from nine BS and 13 HF PBMC samples from the above-mentioned farms to be assessed for the presence of this SNP. Additionally, 17 Sahiwal and 17 Boran cDNA samples were kindly provided by Drs Thomas Tzelos and Tim Connelley (both The Roslin Institute, University of Edinburgh, UK) for TLR2 SNP analysis.

### PBMC Isolation and Macrophage Maturation

Blood for peripheral blood mononuclear cells (PBMC) isolation and subsequent macrophage (MØ) generation was collected from clinically healthy pedigree HF and BS cows. All procedures were carried out under the Home Office license (PPL7009059) approved by the RVC’s AWERB Committee. Blood was drawn into sterile glass vacuum bottles containing 10% v/v acid citrate and centrifuged to separate the buffy coat, which was then diluted in phosphate buffered saline (PBS, Thermo Fisher Scientific, UK) with 2% foetal calf serum (FCS, Sigma-Aldrich, UK). Lymphoprep (*d* = 1.077 g mL^−1^, StemCell Technologies ™, Canada) was underlaid and PBMC were isolated by density gradient centrifugation. PBMC were washed and red blood cells lysed, resultant cells were cultured in RPMI 1640 with GlutaMAX™ (Gibco, UK), supplemented with FCS (Sigma-Aldrich, UK) and penicillin/streptomycin (Thermo Fisher Scientific, UK) at 37 °C with 5% CO_2_. To derive MØ, PBMC were incubated additionally with 10% filtered L929 supernatant. Media was replaced after three days and adherent cells were harvested after 6 days. Matured MØ were phenotypically assessed by flow cytometry using antibodies specific for bovine cell surface expressed markers including CD14, CD16, CD32, CD11b, CD80, CD163 and MHC class II. All antibodies and isotype controls were purchased from Bio-Rad, UK (Table 1). Data were acquired using a FACS Calibur (BD Biosciences, UK) with Cell Quest Pro software (BD Biosciences, UK), counting 10,000 events. Data was exported as FSC files and analysed using FlowJo V10 (FlowJo LLC, USA).

**Table 1.** Antibodies used in this study

| **Antibody** | **Clone** | **Label** | **Dilution** |
| --- | --- | --- | --- |
| **Mouse IgG1 negative control** | - | FITC | 1:20 |
| **Mouse IgG1 negative control** | - | RPE | 1:20 |
| **Mouse IgG2a negative control** | - | FITC | 1:20 |
| **Mouse IgG2b negative control** | - | RPE | 1:20 |
| **Mouse IgG2a anti Human CD 14** | TÜK4 | RPE | - |
| **Mouse IgG2a anti Human CD 16** | KD1 | FITC | - |
| **Mouse IgG2b anti Bovine CD 11b** | CC126 | FITC | - |
| **Mouse IgG1 anti Bovine CD 32** | CCG36 | RPE | - |
| **Mouse IgG1 anti Bovine CD 80** | IL-A159 | FITC | - |
| **Mouse IgG1 anti Human CD 163** | EDHu-1 | RPE | - |
| **Mouse IgG1 anti Bovine MHC II: DR** | CC108 | RPE | 1:25 |
| **Human IgG anti Bovine CD282** |  | AF647 |  |
| **Mouse IgG anti human CD282** |  | AF647 |  |
| **HuCAL Fab-dHLX-MH** |  | AF647 |  |

### RNA extraction and cDNA synthesis

For sequencing and cloning of the *tlr2* gene, total RNA was isolated from bovine PBMC using a RNeasy Mini kit (Qiagen, UK) with on-column gDNA digestion (DNAse from Ambion, UK) according to the manufacturer’s instructions. Total RNA was reverse transcribed to cDNA using iScript cDNA Synthesis Kit (Bio-Rad, UK) according to the manufacturer’s instructions. cDNA samples were checked for concentration and purity via the A260/280 ratio using a Nanodrop ND-1000 Spectrophotometer (Thermo Fisher Scientific, UK). A RT-negative control of each RNA sample was analysed by polymerase chain reaction (PCR) for expression of β-actin to confirm lack of gDNA contamination of RNA.

### Cloning and sequencing of *tlr2*

Full length *tlr2* sequences were amplified using primers designed using Primer-BLAST (NCBI, USA) and Primer3Plus (Table 2) (187) based on bovine genome Bos_taurus_UMD_3.1.1 (NCBI, USA) as reference. PCR reactions were performed using Easy-A High-Fidelity PCR master mix (Agilent Technologies, UK) on a Mastercycler Pro Thermal Cycler (Eppendorf, UK). Following amplification, the PCR products were purified using the MinElute PCR purification kit (Qiagen, UK) according to the manufacturer’s protocol. For TA cloning of full length bovine *tlr2* into the pDrive plasmid (Qiagen, UK) or pGEM-T plasmid (Promega, UK), manufacturer’s instructions were followed. Plasmids were transformed into NEB^®^ 10-beta competent *E. coli* (New England Biolabs, UK) and colony PCR performed to verify the presence of the *tlr2* insert. Sequences were validated using Sanger sequencing; DNA sequencing was performed by DNA Sequencing & Services (MRC I PPU, School of Life Sciences, University of Dundee, Scotland, www.dnaseq.co.uk) using the corresponding vector-specific sequencing primers. A minimum of 4 different plasmids derived from unique colonies were sequenced for each animal. Bovine TLR2 contigs were assembled using the bovine TLR2 sequence *Bos taurus* Hereford breed NM_174197.2 (NCBI, USA) as a reference. TLR2 nucleotide sequences were translated into corresponding amino acid sequences and aligned in CLC Main Workbench V7.6.4 (CLCbio, Denmark). Detected SNPs were classified according to the dbSNP database (NCBI, USA). Analysis of SNP data was curated and as at present, the NCBI dbSNP database contains only human data, allele changes, CDS position and function of each SNP was manually cross-checked and referenced against Ensemble and the European Variation Archive (EVA).

**Table 2.**
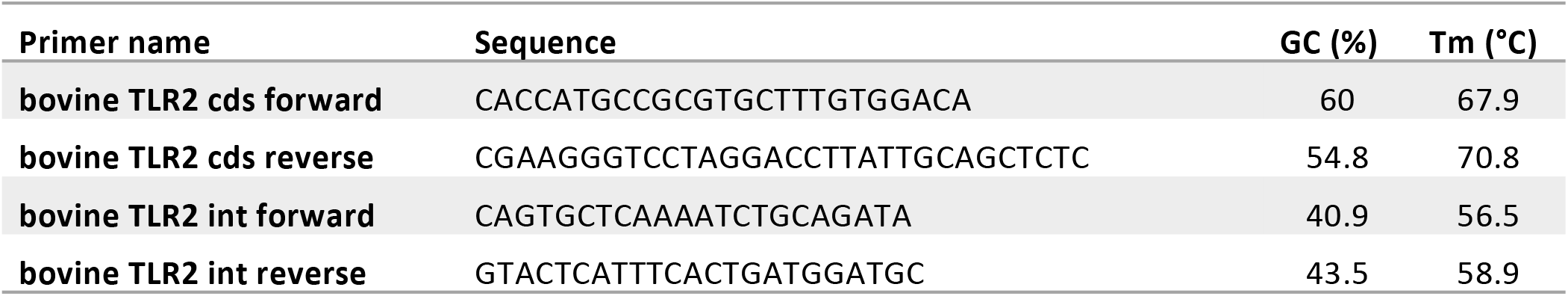
Primer names for PCR reactions. All primers were designed using Primer-BLAST (NCBI) and Primer3Plus (http://www.bioinformatics.nl/cgi-bin/primer3plus/primer3plus.cgi)

### Expression of TLR2 by HEK 293T (SEAP) cells

The Human Embryonic Kidney (HEK) 293T cell line, stably transfected with the gene encoding NF-κB inducible secreted embryonic alkaline phosphatase (SEAP), was cultured SEAP media as described recently [54]. Bovine or human TLR2 sequences of interest were cloned into pcDNA3.3 TOPO TA mammalian expression vector (Thermo Fisher Scientific, UK), and transfected into SEAP HEK cells using TurboFect transfection reagent (Thermo Fisher Scientific, UK) according to the manufacturer’s protocol. After 24 h, cells were counted and re-seeded in triplicates in a 96-well plate to rest for another 24 h before either evaluation of transfection efficiency or stimulation with various agonists. Surface expression of TLR2 was confirmed by flow cytometry using either human anti-bovine CD282 (anti-TLR2) antibody or mouse anti-human CD282 antibody, both conjugated with Alexa Flour 647 (Bio-Rad, USA). HuCAL Fab-dHLX-MH Ab (Bio-Rad, USA) was used as a negative control as recommended by the manufacturer.

### Stimulation Assays

SEAP HEK cells expressing TLR2 constructs, or PBMC-derived bovine MØ were specifically stimulated using 100 ng mL^−1^ diacylated FSL-1 (Invivogen, USA); 1 mg mL^−1^ of Pam_3_CSK_4_; 19 kDa lipoprotein antigen (represents Rv3763 or Mb3789) (EMC microcollections, Germany), M. *bovis* BCG (MOI of 10) or MTB H37Rv at (MOI of 5). The direct NF-κB activator phorbol 12-myristate 13-acetate (PMA, Sigma-Aldrich, UK) was used at 200 ng mL^−1^ as a positive control. Sterile culture media and mock transfection with an empty vector plasmid were used as negative controls. Supernatants were harvested for CXCL8 ELISA as described by Cronin *et al.* (185) for bovine MØ, or Quantikine Human CXCL8 ELISA kit (R&D Systems, USA) for human MØ, or colorimetric SEAP assay from HEK cells. SEAP activity as induced by NF-kB activation was assessed by the addition of pre-warmed Quantiblue reagent (Invivogen, USA) followed by incubation for 24 h at 37°C. For both assays, optical density was measured using a SpectraMax M2 spectrometer (Molecular Devices, UK).

## Results

### Brown Swiss genome assembly and QC

We used the 10x Genomics tool Supernova to assemble a haplotype-phased Brown Swiss cow genome. After trimming, the mean read length was 138.5 bases, with a raw read coverage of 59.64. The genome size was calculated to be 2.66Gb (of which 98.08% of the genome was present in scaffolds >10Kb) with a contig N50 of 523Kb and a scaffold N50 of 26Mb (**Supplementary Table 1**). We used BUSCO to identify reconstructed single copy orthologs. From 9226 BUSCO orthologs 95.2% were recovered, 93.1% were single-copy, 2.1% were duplicated, 1.5% were fragmented, and 3.3% were missing. We used KAT to examine the distribution of kmers in the assembly to that in the reads (Error! Reference source not found.). The genome shows a normal distribution of kmers found once in the assembly and once in the reads (red distribution). It also shows a number of low frequency kmers (black peak between 0-10) present in the reads, but not in the assembly that most likely represent sequencing errors removed from the assembly. This continues into a shallow distribution of kmers missing from the assembly (black distribution between 10-40), which most likely represents haplotypes found only in the alternate phased assembly.

### Brown Swiss genome annotation and gene expansion

We noted genes that were missing **(Supplementary Table 2)** and duplicated in the Brown Swiss genome, ordering the duplicated genes in order of the most numerous duplications (**Supplementary Table 2**). We note that all genes annotated as having five or more copies belonged to one of five genes. Three of these genes were repeat/transposon related genes. The remaining two genes were a predicted Bos indicus x Bos taurus elongation factor 1-alpha 1 pseudogene (6 copies), and the Bos taurus T cell receptor gamma cluster 1 (TCRG1) gene (128 copies).

### Identification of single nucleotide polymorphisms (SNPs) within bovine *tlr2* between cattle breeds

TLR2 is an important PRR involved in the recognition and uptake of MTBC family members. Key SNPs in TLR2 have been identified that affect susceptibility to mycobacterial infection for both humans and cattle [55,56]. The CDS of the bovine *tlr2* gene of six BS and four HF cattle were cloned and compared to the bovine NCBI reference sequence genome (RefSeq) comprised of that of a *Bos taurus* Hereford breed bull (NM_174197.2, NCBI, USA, accessed April 2017) in order to identify potential functional SNP sites. A total of 19 SNP variants were detected across the breeds, comprising both synonymous and missense mutations (**Table 3**). Of the eight missense mutations, seven occur in the ECD region of TLR2 and one in the TIR domain. One synonymous SNP at mRNA position 202 could not be classified with any available database, was described by Jann et al [47]and is potentially novel. Of all SNPs identified across the breeds, two of the non-synonymous SNPs have been already reported in the literature [44,47,55] and were only observed in the BS animals. One identified SNP variant, rs55617172, leading to an amino acid change of aspartic acid (D) to glutamic acid (E), both negatively charged, at amino acid position 63, has been significantly associated with TB resistance in a TB case-control cattle study [44]. However, SNP rs68343167, was only identified in BS isolates in this study, and results in an amino acid change of the positively charged histidine (H) to the uncharged glutamine (Q) at amino acid position 326 (H326Q). This amino acid position has been suggested to be of relevance for ligand binding and functionality of human TLR2 [47,57], bovine TLR2 and was additionally identified in the Anatolian Black *Bos taurus* breed [55]. In addition, position 326 resides in leucine rich repeat 11 (LRR11) which together with LLR12 are the key domains involved in ligand biding and heterodimerisation of TLR2 with TLR1 and TLR6 [57].

**Table 3:**
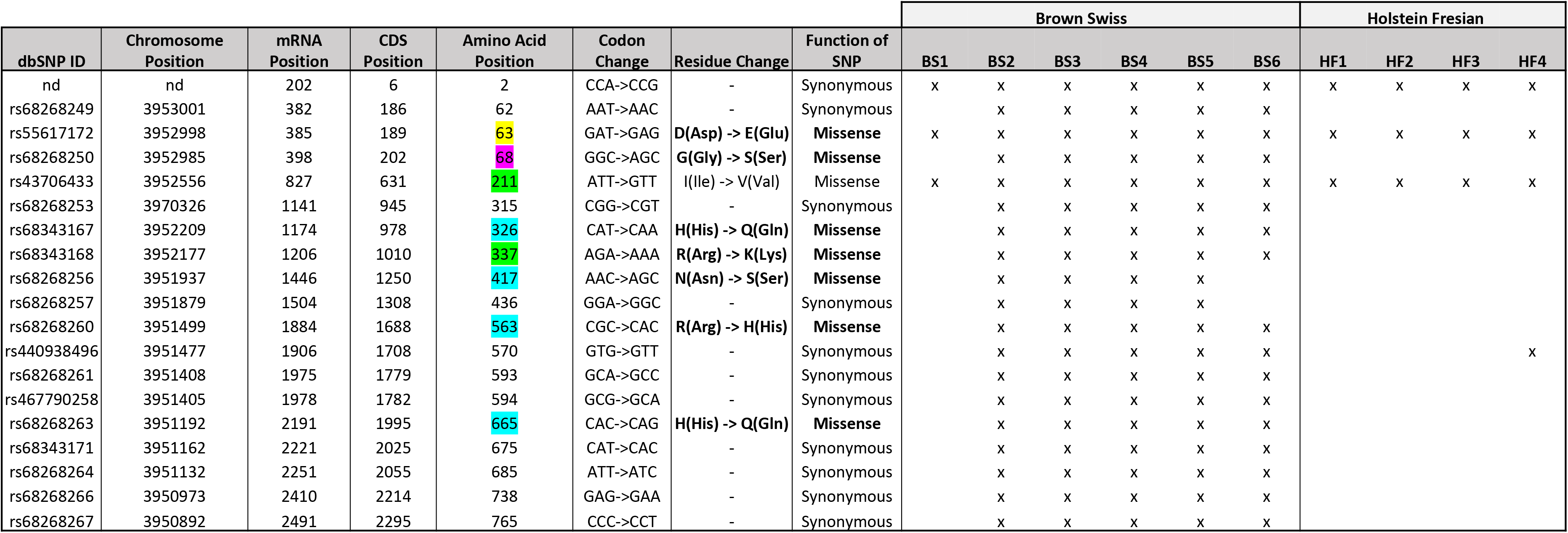
Identified SNP variants within the coding sequence of bovine *tlr2* in HF and BS breeds animals. The coding sequence for TLR2 in BS (n=6) and HF (n=4) animals were identified by Sanger sequencing and contigs matched against reference sequence NM_174197.2 (*Bos taurus* Hereford breed). SNP variants were classified as either synonymous or non-synonymous mutations, with the corresponding amino acid changes displayed. The SNP detected at mRNA position 202 was not found in the NCBI dbSNP database. Presence of allele change within individual animals is depicted as ‘x’. Amino Acid position highlighted depicts positions noted from TLR2 CDS transcript alignments detailed in Supplementary Figure S4. Yellow = unique to BS, Green = unique to BS and *B. taurus*, Blue = unique to *B. indicus* and Magenta = variable across all species.

### Genotype frequency analysis of candidate SNP H326Q in selected cattle breeds

Having identified multiple SNPs within the coding sequence of *tlr2* gene across breeds, SNP rs68343167 (H326Q) was selected for further investigation using a larger sample size, including cDNA from two additional breeds, Boran and Sahiwal, from the *B. indicus* species. A total of 66 cDNA samples (Brown Swiss n=15, HF n=17, Sahiwal n=17, Boran n=17) were assessed for H326Q SNP analysis and frequency analysis (**Table 4**). The reference codon at the site of interest is encoded by the CAT sequence (TT genotype). Individual animals carrying the SNP had either heterozygote codons (TA genotype), encoded by either CAT or CAA, which is responsible for the H326Q variant; or they were homozygote for the CAA sequence (AA genotype).

**Table 4:**
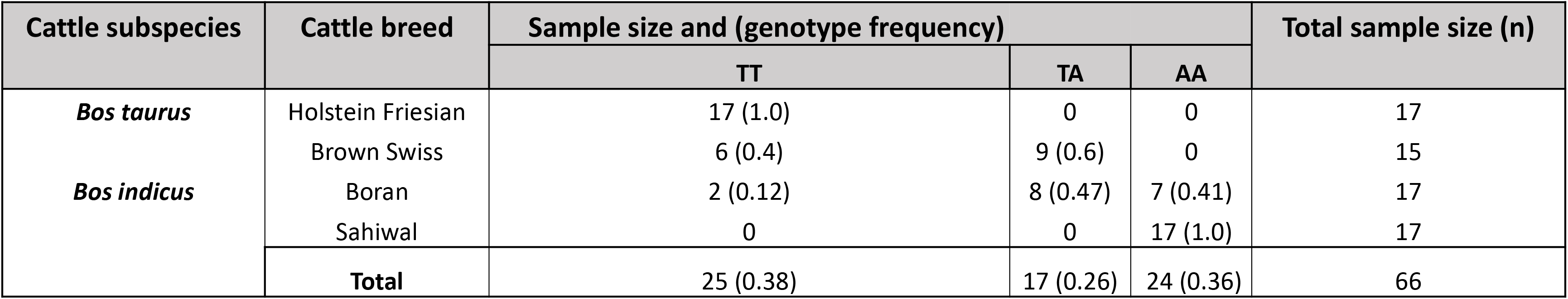
Genotype frequency of *Bos taurus* and *Bos indicus* cattle breeds for TLR2 selected candidate SNP rs68343167 (H326Q). A total of 66 cattle from *Bos taurus* and *Bos indicus* breeds (BS n=15, HF n=17, Sahiwal n=17, Boran n=17) were examined for their genotype frequencies with respect to SNP rs68343167 (H326Q). Genotype ‘TT’ codes for the homozygous ‘wildtype’ bovine TLR2 reference sequence NM_174197.2. Genotype ‘TA’ were heterozygous individuals and genotype ‘AA’ were homozygous for the H326Q SNP variant.

Interestingly, all HF samples tested as part of the present study were homozygous for the *B. taurus* Hereford RefSeq (TT genotype). Of the BS samples, six samples were homozygous for the RefSeq sequence, whereas the majority (n=9) were found to be heterozygous (TA genotype), containing both, the RefSeq sequence and the H326Q SNP. Of the *Bos indicus* breeds, the Boran samples revealed two samples homozygous for the RefSeq sequence, eight heterozygous and seven homozygous for the SNP. Thus, the heterozygote genotype was most abundant in the BS and Boran population. All Sahiwal samples (n=17) were homozygous for the SNP variant (AA genotype).

Alignment of TLR2 transcripts revealed additional SNP sites unique to BS, BS and *B. taurus* as well as *B. indicus*. Of note are positions H326Q as reported here and by others, as well as positions 417, 563 and 665; these latter 3 amino acid positions are particularly interesting as like H326Q these SNPs were found to introduce identical residue changes to our full length TLR2 analysis. Further, these SNPs were found to be present in BS cattle investigated (Table 3) but not present for HF cattle suggesting a role for genotype impact as found for H326Q. Position 417 and 665 are noteworthy, belonging to LRR15 and the TIR domain respectively. As LRR15 is close to the LRR regions thought to be involved in dimerization of TLR2 with co-receptors this SNP may play a role in effective PRR dimers forming. As the TIR domain is crucial for receptor dimerization of TLR cytoplasmic domains as well as signal potentiation; SNP H665Q may also modulate functional responses of TLR2.

On establishing distinct genotypes for the H326Q SNP in a range of cattle species (i.e., HF = TT, BS and Boran = TA and Sahiwal = AA), we assessed in the next step this association with TLR2 gene models and global sequence alignments. From the genotype analysis BS and Boran cattle represent an intermediate between the genotype extremes, perhaps representing genomic heritage; for example, Boran cattle are predominantly *B. indicus* with underlying influences of European and African *B. taurus* [58,59]. TLR2 gene models comparing the BS genome to *B. taurus*, *B. indicus* and *B. bubalis* (water buffalo) highlights an alignment of BS with *B. taurus* as TLR2 is derived from a single transcript in both breeds (Supplementary Figure S3). Highlighting the heterozygous genotype of BS cattle, BS resembled *B. taurus* at H326Q within LRR11 of the extracellular domain of TLR2 (Supplementary Figure S4) when the CDS of each possible TLR2 transcript was aligned to the three cattle species, Interestingly, additional sites where the BS and *B. taurus* sequences differed from *B. indicus* where I211V, F227L and R337K. Positions 211 and 337 were identified from the initial TLR2 sequencing described above.

### The Brown Swiss H326Q TLR2 polymorphism induces a stronger NF-κB response after ligand binding compared with Hereford and Holstein Frisian TLR2

Having identified the H326Q candidate SNP and its representation in the present study population of different cattle breeds, specifically in the BS population, the potential impact of this SNP on boTLR2 function in this breed was assessed. We utilised a previously published HEK cell assay to express the different TLR2 constructs in the absence of additional PRR [54]. As well as the BS H326Q TLR2 variant, we included the Hereford reference variant and a L306P SNP variant that has been identified by others [47,55] but not identified in the present study. We were also interested to assess the functionality of TLR2 across species, and therefore included the huTLR2 reference sequence (NCBI RefSeq Accession Number NG_016229.1). Furthermore, novel chimeric TLR2 constructs featuring the ECD of huTLR2, with the TIR domain of boTLR2 were created to delineate the contribution of each component to TLR2 functionality.

To confirm that the different TLR2 constructs were similarly expressed on the cell surface of the SEAP HEK cells, immunostaining was performed using bovine and human TLR2 specific antibodies. The mean fluorescence intensity (MFI) values of the positively stained population were similar for all constructs (**Figure S1**). Stimulation with PMA, a non-specific activator of NF-κB, was used as a positive control and resulted in consistently high SEAP secretion for all transfectants. SEAP activity in response to ligands was normalised to PMA responses as an internal control.

The H326Q variant showed significantly higher TLR2-dependent NF-кB response in comparison to RefSeq Hereford TLR2 (p < 0.001 **Figure 2 A and C**). Constructs containing the L306P SNP did not respond significantly differently in comparison to the RefSeq Hereford boTLR2 version. The response of SEAP HEK cells transfected with huTLR2 was significantly higher in response to both FSL-1 and Pam_3_CSK_4_ in comparison to the bovine constructs. Interestingly, the chimeric bov:hu TLR2 receptor showed higher NF-κB activity than the Hereford boTLR2 HEK cell line, but significantly lower activity than the huTLR2 receptor in response to both FSL-1 and Pam_3_CSK_4_.

### NF-κB activity correlates directly to CXCL8 production of TLR2-HEK cells upon stimulation with lipopeptides

Using cell culture supernatant recovered from the above SEAP assay experiments, CXCL8 production in response to FSL-1 and Pam_3_CSK_4_ stimulation were measured (**Figure 1 B and D**). As previously, there was an increase in CXCL8 production in the H326Q variant in comparison to the bovine RefSeq Hereford TLR2, however this was found only to be statistically significant in response to FSL-1 stimulation (p<0.01). CXCL8 production measured in huTLR2 transfectants was significantly higher than in boTLR2 transfectants in response to both ligands. CXCL8 production by chimeric bo:hu TLR2 HEK cells was significantly higher than any of the boTLR2 variants to FSL-1 (p<0.05), but lower than those from huTLR2, reflecting the observations made using the SEAP reporter assay of NF-κB activity.

**Figure 1:**
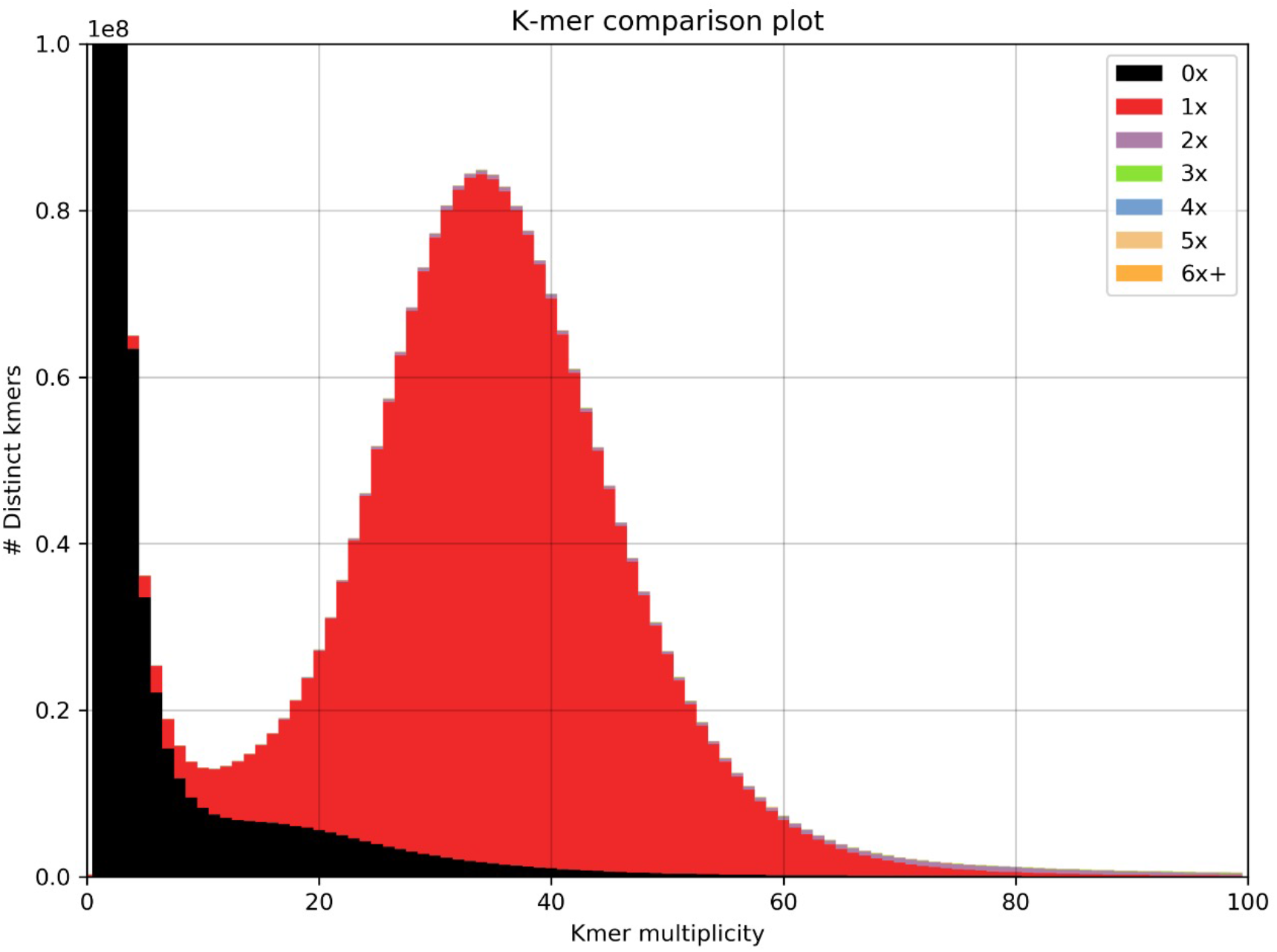
Distribution of kmers. One haplotype of the Brown Swiss genome compared to that of the complete read set (minus 10x barcodes). The red distribution signifies the distribution of read-kmers found only once in the assembly and the black distribution signifies the distribution of read-kmers not found in the assembly (mainly sequencing errors and haplotypes present in the alternate phased assembly).

### huTLR2 is significantly more sensitive than boTLR2 variants to stimulation with live *M. bovis* BCG, but not *M. tuberculosis*

Having assessed the activity of TLR2 variant HEK cells using TLR2-specific agonists and determining good correlation between SEAP activity and CXCL8 secretion (**Supplementary Figure S3**), we next challenged the TLR2-HEK cells with live mycobacteria and measured the NFkB activation responses. Interestingly, the response of SEAP HEK cells expressing huTLR2 to challenge with *M. tuberculosis* was markedly lower than when they were challenged with TLR2-specific lipopeptides i.e. Pam_3_CSK_4_ and FSL-1 in the previous experiment (**Figure 2 and Figure 3**). Only the huTLR2 variant produced a significant NFkB response upon challenge with *M. bovis* BCG compared to mock simulations. NFkB responses for H326Q containing TLR2 were significantly lower than Hereford reference TLR2 and those constructs containing L306P. All boTLR2 variants responded more strongly to *M. tuberculosis* than to *M. bovis* BCG, even though the MOI was lower (MOI of 5) than with the *M. bovis* BCG infection (MOI of 10).

**Figure 2:**
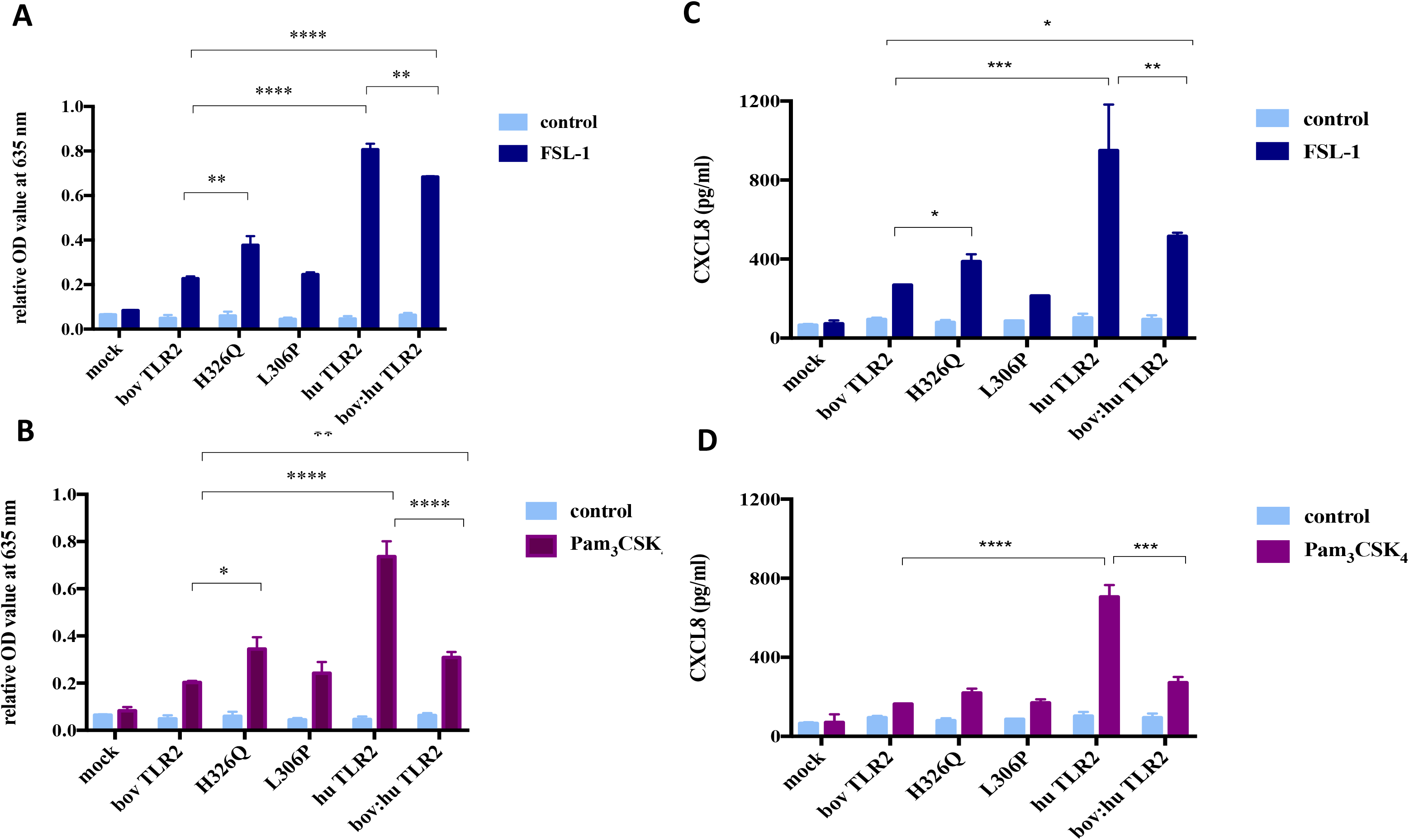
TLR2-Ligand-Dependent NF-κB Activity and CXCL8 Secretion by HEK 293 cells Expressing Bovine and Human TLR2 Sequence Variants. HEK 293 cells, harbouring NF-κB-induced SEAP reporter genes were transfected with either empty pTracer vector (mock); bovine or human TLR2 CDS constructs, or a bovine: human chimera in pDuo-mcs. Cells were stimulated with either FSL-1 at 100 ng/ml (A,C), Pam3CSK4 at 1 mg/ml (B,D) for 24 hrs. SEAP activity was measured by quantifying optical density of cell culture supernatant at 635 nm (A,B). OD values were normalised against values for PMA stimulation at 200 ng/ml. Additionally, supernatants were assessed for CXCL8 concentration by ELISA (C,D). Error bars represent standard deviation from the mean of triplicate technical replicates and are representative of three independent repeats. (*=p<0.05, **=p<0.01, ***=p<0.001, ****=p<0.0001) and are only shown in relation to bovine TLR2 and additionally between the human and the chimeric bov: hu TLR 2 receptor. All data analysed by two-way ANOVA with multiple comparisons using GraphPad Prism V8 (GraphPad Inc., USA).

**Figure 3:**
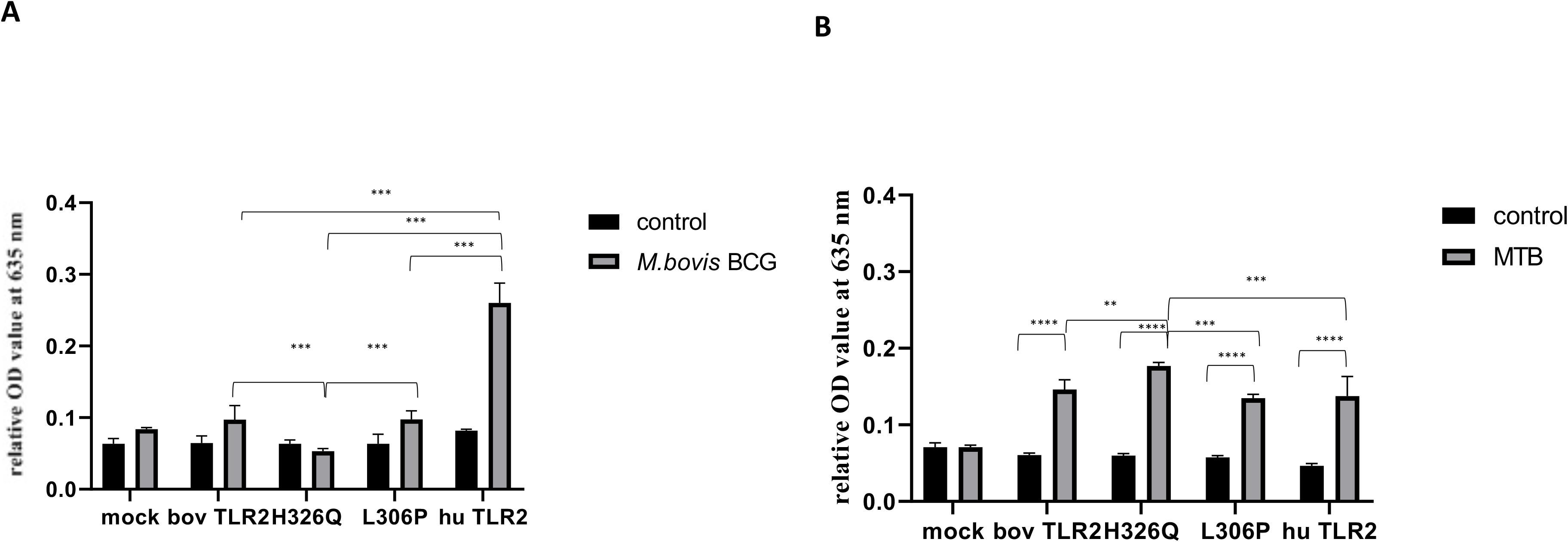
*M. bovis* BCG and *M. tuberculosis*-Dependent NF-κB Activity Secretion by HEK 293 cells Expressing Bovine and Human TLR2 Sequence Variants. HEK 293 cells, harbouring NF-κB-induced SEAP reporter genes were transfected with either empty pTracer vector (mock); bovine or human TLR2 CDS constructs, or a bovine: human chimera in pDUO-mcs. Cells were stimulated with either live *M. bovis* BCG (MOI of 10) (**A**) or *M. tuberculosis* H37Rv (MOI of 5) (**B**) for 24 h. SEAP activity was measured by quantifying optical density of cell culture supernatant at 635 nm. OD values were normalised against values for PMA stimulation at 200 ng/ml. Error bars represent standard deviation from the mean of triplicate technical replicates and are representative of three independent repeats. (*=p<0.05, **=p<0.01, ***=p<0.001, ****=p<0.0001) and are only shown in relation to bovine TLR2 and additionally between the human and the chimeric bov: hu TLR 2 receptor. All data analysed by two-way ANOVA with multiple comparisons using GraphPad Prism V8 (GraphPad Inc., USA).

### Brown Swiss and Holstein Friesian PBMC-derived macrophages respond significantly differently to TLR2-specific and mycobacterial ligands

Differences in functionality at the whole cell level conferred by boTLR2 breed variants were further assessed using PBMC-derived MØ. Cells were isolated from BS, containing the heterozygote H326Q variant, or HF cattle, which have the TLR2 sequence identical to the RefSeq Hereford breed. Due to restriction with class 3 lab access, we measured initially the CXCL8 response to the TLR2-specfic agonist, FSL-1 as well as to BCG. In agreement with TLR2-HEK cell responses (**Figure 2**), MØ generated from BS carrying the H326Q variant showed a significantly higher CXCL8 responses than HF MØ upon FSL-1 stimulation (p<0.05, **Fig 4A**). We further verified a high degree of correlation between the SEAP NF-kB and CXCL8 assays in a control experiment, by collecting data for both from the same stimulated cells and performing regression analysis (Fig S2). Interestingly, unlike the SEAP HEK cell experiments, the PBMC-derived MØ responded similarly to MTB-derived 19 kDa lipoprotein and *M. bovis* BCG compared with FSL-1. We also detected a significantly greater response by BS derived MØ to *M. bovis* BCG compared to untreated cells, while the response by HF-derived MØ was not statistically greater than untreated cells.

**Figure 4:**
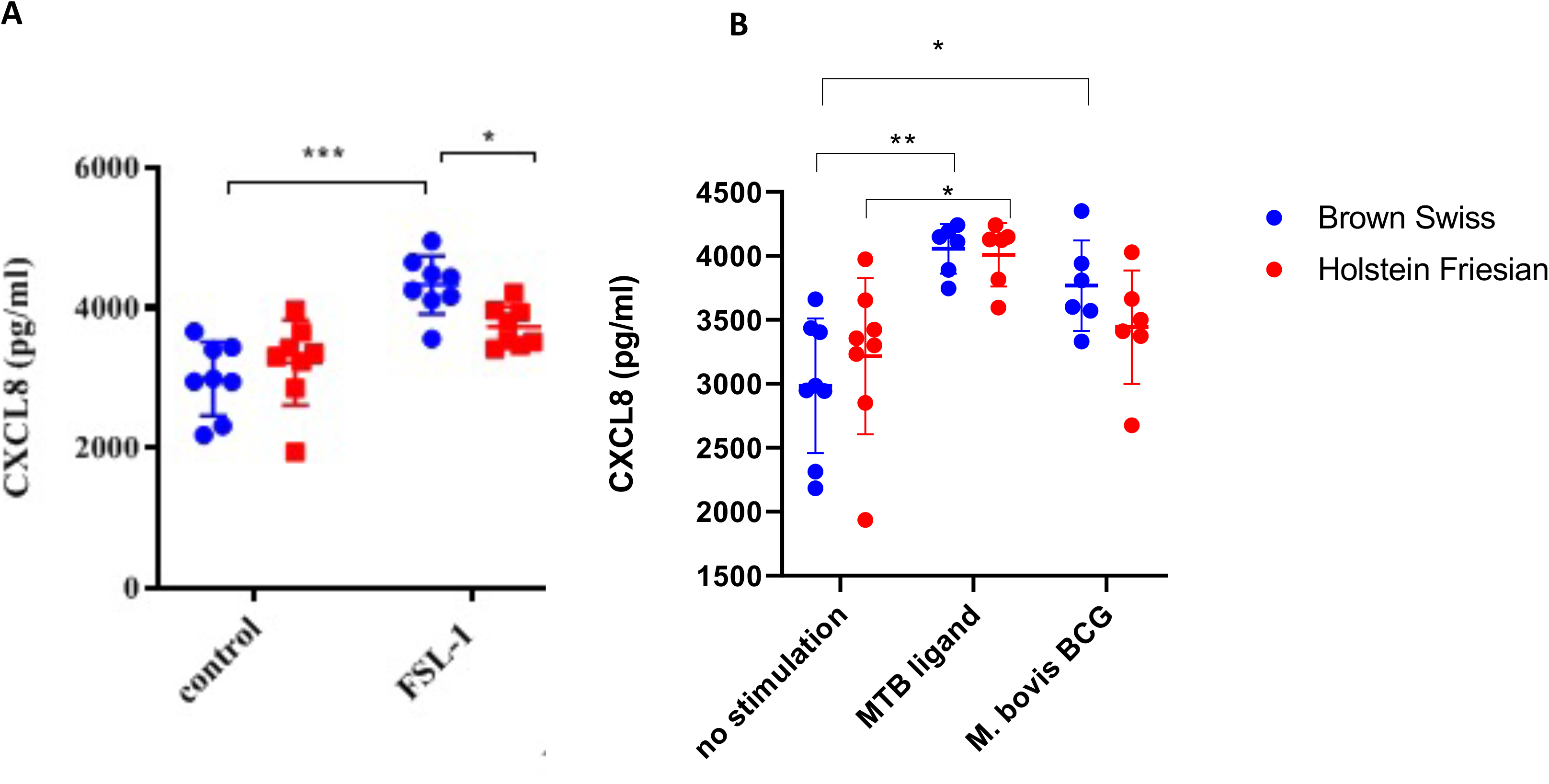
CXCL8 production by bovine MØ in response to FSL-1, *M. bovis* BCG or MTB Ligand Stimulation. Bovine MØ of BS (heterozygous for TLR2 H326Q SNP) and HF MØ (homozygous for the wild-type TLR2 sequence) were stimulated with either FSL-1 at 100 ng/ml (**A**), 19 kDa lipoprotein antigen (represents Rv3763 or Mb3789) (EMC microcollections, Germany) or infected with *M. bovis* BCG at MOI = 5 (**B**) for 24 hr. CXCL8 concentration in cell culture supernatant was assessed by ELISA. A total of n=8 animals per breed were stimulated with FSL-1 and n=6 animals per breed were infected with *M. bovis* BCG. Error bars represent standard deviation from the mean. Data was analysed with two-way ANOVA and Tukey’s post hoc comparison in GraphPad Prism V8 (GraphPad Inc., USA) (*=p<0.05, ***=p<0.001).

## Discussion

Several studies have investigated cattle breed resistance to various diseases at the genetic level, leading to the suggestion that the HF breed may have lost immunological fitness, possibly due to extensive breeding for production traits [56,60]. Recently, we have also reported breed-specific differences in response to mycobacterial challenge (60). It is estimated that the cost of bovine tuberculosis to UK taxpayer is in excess of £100 million per year in surveillance testing and compensation, and there is currently no viable vaccine for cattle. Investigation into aspects of host immunity as they relate to different cattle breeds may inform future breeding strategies and minimise the impact of disease and reduce transmission of zoonotic TB to humans. Zoonotic TB is currently estimated to account for ~10% of global TB cases therefore taking a One Health approach to tackle zoonotic TB is paramount in achieving the UN Sustainable Development Goal of ending the TB epidemic [61]. Reducing the burden of *M. bovis* infection in animal reservoirs such as livestock and wildlife underpins the 10 priority areas laid out in the WHO Roadmap for Zoonotic TB [61].

Using the power of linked-read sequencing, we have generated the first *de novo* phased assembly of the Brown Swiss cow genome. The genome shows high contiguity and completeness, as shown by the relatively high scaffold N50 and high single-copy ortholog reconstruction. Genome annotation lift over and gene duplication analyses show an increase in gene copy number of T cell receptor gamma cluster 1 (TCRG1) genes, in line with cattle belonging to “high γδ species” compared to human counterparts [62–64]. It is important to note is that γδ T cells are not only considered a major group of cells in mucosal immune responses, but also have been shown to be important in early immune responses to mycobacterial infection and bridge the gap between innate and adaptive immunity [65].

We identified 19 SNPs occurring within the CDS of bovine TLR2 compared to the RefSeq. A H326Q mutation was of particular interest as it resulted in an amino acid change from the positively charged histidine (H) to the uncharged glutamine (Q) at position 326, which is located in the ligand binding region LRR11 of the ECD of the receptor [57]. Using a phylogeny-based approach, this SNP has been previously identified as a mutation under positive selective pressure in cattle and suggested to be of functional relevance [47]. Furthermore, the H326Q variant was found by Bilgen *et al*. [55] in the *Bos taurus* Anatolian Black cattle breed, which is thought to be more resistant to pathogens recognized by TLR2, such as to *M. bovis* and mastitis-causing bacteria. Among the *Bos indicus* breeds, all Sahiwal samples in our study were homozygous for the H326Q SNP and except for two samples, half of the Boran samples were either homozygous or heterozygous for this variant, confirming a strong presence of this SNP in the *Bos indicus* samples tested. This is interesting given that these breeds are known to be less extensively bred for production traits [55] and more resistant to several diseases, including bovine tuberculosis [56] compared with *Bos taurus*. Within the BS breed, more than half were heterozygous for the SNP. For HF samples, the cattle breed under strongest selective pressure for production traits in our study population, all samples were homozygote and identical to the reference Hereford sequence. Since *Bos indicus* cattle are phylogenetically more ancient than *Bos taurus* cattle with respect to their common ancestor *Bos primigenius* [66], our results suggest that BS cattle are immunologically potentially more closely related to ancient cattle breeds such as Boran and Sahiwal than the HF breed.

To compare the functional relevance of the TLR2 variants present in the HF and BS breeds, the identified sequences were cloned and expressed in HEK cells for *in vitro* phenotypic assessment. Initially, we used the SEAP assay that has been previously reported as a quantitative indicator of NF-κB activation during TLR stimulation [54,67] and in which the SEAP response was directly correlated to CXCL8 production (22, 61). Upon stimulation with synthetic lipoproteins and BCG, constructs containing the H326Q SNP, showed a significant increase in NF-κB activity and therefore TLR2 signalling strength. As the H326Q SNP was well represented in our *Bos indicus* population and most BS samples were heterozygous for this SNP, it can be hypothesized that this primes for a stronger TLR2-dependent immune response among these cattle.

For comparison, we included the characterisation of another reported SNP in the ECD domain of boTLR2, the L306P variant [47,57] – although this was not identified in the present study. While this site is reported to be important for determining the size of the ligand-binding groove [57], we did not detect significantly different NFkB or CXCL8 response to stimulation with TLR2-specific ligands used in this study. A downstream functional outcome of NFkB activation is the transcription of certain inflammatory cytokines and including CXCL8 [68,69], which is responsible for the recruitment of neutrophils to the site of infection and thus an important regulator of innate immunity. We observed a high degree of correlation between NFkB activity and CXCL8 production in our assay, confirming the functional relevance of the SNPs between cattle breeds.

TLR function has been extensively studied, and shown to be species-specific and dependent on the ECD of the receptor [70]. In contrast, there are conflicting reports over the contribution of the TIR domain, which is more highly conserved between species [7,9] [70–72]. We sought to clarify this aspect in the context of TLR2 expressed in isolation and challenged with TLR2-specific ligands. For both FSL-1 and Pam3CSK4, the bovine TIR showed effective, but reduced activity when expressed as a chimera with the huTLR2 ECD compared with the native human ECD-TIR structure. Similarly, the human ECD imparted greater TLR2 activity as a chimera with bovine TIR compared with native bovine ECD-TIR structure. Therefore, we can confirm that both ECD and TIR are independently responsible for differences in TLR2 sensitivity between species when TLR2 is expressed as a homodimer. In our study the human variant of both is more sensitive than the bovine orthologue to TLR2 ligands in the absence of other PRR. This contrasts with findings previously reported where the ECD was responsible for species-specific responses when a co-receptor construct of TLR2-Dectin1 was used [73], however these interactions may highlight the importance of co-receptor signal potentiation to increase the repertoire of innate immune responses. Our findings support the work of Faber et al who demonstrated a similar result with human and porcine TLR5 chimeras [71].

Interestingly, the magnitude of the TLR2-dependent response was reduced when the cells were stimulated with live mycobacteria than with the synthetic ligands, independent of SNPs. There were some statistical differences between SNPs in NFkB activity, but none of these were significantly different from the mock untreated conditions and so are not likely to be biologically relevant. *Mycobacterium* spp. have been reported to be capable of inhibiting the TLR2-driven response - for example, by the expression of glycolipids such as PDIM in the cell envelope that can obscure TLR2-ligand recognition [74]. Furthermore, some MTB PAMPs may optimally require TLR2 heterodimerization or engagement of additional PRRs such as CLRs and DC-SIGN to interact and induce NFκB [14,75,76]. By contrast, FSL-1 and Pam_3_CSK_4_ [77] have been shown to activate TLR2 as a homodimer [54] or TLR 2/6 heterodimers [78].

The response of SEAP HEK cells expressing bovine TLR2 was significantly higher when challenged with *M. tuberculosis* than with *M. bovis* BCG. Conversely, cells expressing huTLR2 were significantly more sensitive to *M. bovis* BCG than cells expressing boTLR2. As *M. bovis* BCG is derived from *M. bovis*, this may reflect species-level differences in host-pathogen coevolution that has resulted in humans being generally more resistant to *M. bovis* infection than cattle and *vice-versa*, cattle being more resistant to *M. tuberculosis* [79]. *M. bovis* BCG is the avirulent vaccine strain of *M. bovis* that in certain circumstances induces protective immune but does not induce disease suggesting that modulated responses compared to pathogenic mycobacteria could be expected. It is interesting that this effect was observed when TLR2 was expressed in the absence of additional PRR. While heterodimerization between PRR has been shown to broaden the ligand spectrum for TLR2, it is not thought to induce differential downstream signalling pathways [80]. The presence of accessory molecules such as CD14 and CD36 have been described to enhance TLR2 pathogen recognition by concentrating microbial products on the cell surface [81] and potentially this might be required for efficient boTLR2 recognition of *M. bovis* but is dispensable for recognition of *M. tuberculosis*. Further work is required to delineate the response of both species to virulent *M. bovis* in comparison with *M. bovis* BCG.

Whereas the HEK cell model enabled assessment of TLR2 function in isolation, we further characterised PBMC-derived MØ from either breed to examine a potential role in the context of additional PRR. BS MØ produced significantly higher CXCL8 to the TLR2-specific lipopeptide, FSL-1, than MØ from HF animals. Furthermore, BS macrophages responded significantly better than unstimulated controls in response to *M. bovis* BCG, whereas HF macrophages did not. Taken together, our data are consistent with a positive role for the H326Q TLR2 variant supporting innate immunity at the whole cell level to mycobacterial infection. Another finding was that PBMC-derived MØ responded with greater CXCL8 production than HEK-TLR2 cells. This supports previously published observations that while TLR2 can initiate CXCL8 production, additional pathways can also upregulate the chemokine [11][82]. Therefore, the SNP in TLR2 contributes to, but is not solely responsible for the inflammatory response to mycobacteria. It is likely that additional breed-specific factors, including a possible contribution by adaptive immunity determines the bovine response to tuberculosis at the whole animal level.

Our findings provide proof of concept however, that meaningful genetic differences exist between cattle breeds that impact immune function, and such work can be expanded to consider additional factors. It is also interesting to speculate which of the identified TLR2 sequences may actually represent the “wild-type” sequence, and whether less responsive cells for HF animals are actually hyporesponsive, due to continues selection for production, whereas the sequence and response seen in BS does represent the “norm”. Resistance to disease is often multifactorial and polygenic and thus identified resistance markers are often unlikely to provide absolute resistance to disease [83]. Of course selection for resistance should not occur at the expense of other control measures such as biosecurity and bTB control in the wildlife reservoir [84]. Immune function and mycobacterial resistance factors associated between cattle breeds (and species) should be considered when implementing vaccine-based or immunomodulation-based control measures to reduce *M. bovis* burden.

## Acknowledgements

DNA samples from Boran and Sahiwal cattle where kindly provided by Drs Thomas Tzelos and Dr Tim Connelley, Roslin Institute, Edinburgh, UK.

