## Supplementary Tables and Figures for "Single Nucleotide Polymorphisms in the Bovine TLR2 Extracellular Domain Contribute to Breed and Species-Specific Innate Immune Functionality"

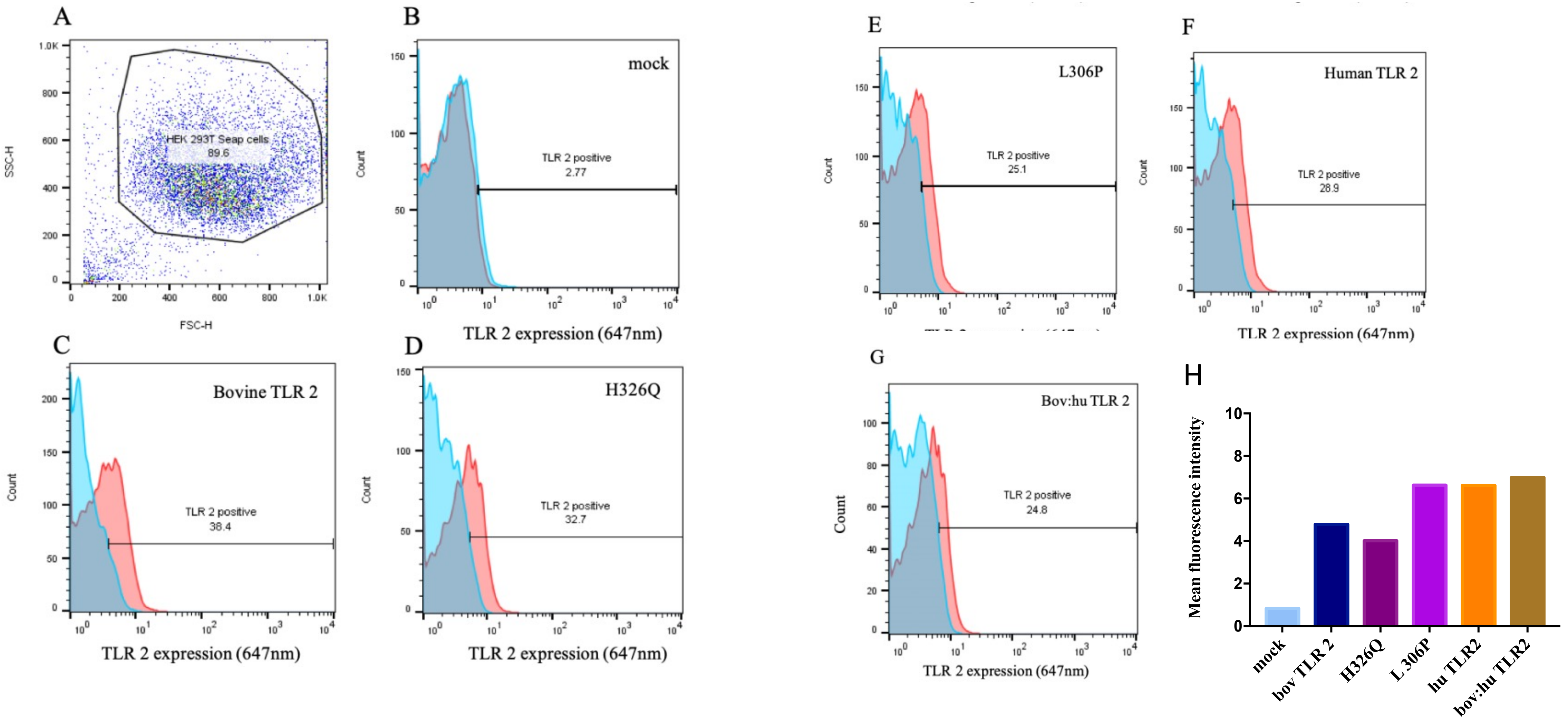


**Figure S1: TLR2 receptor expression in HEK 293 SEAP cells upon transient transfection.** Upon transient transfection with different TLR2 constructs, surface expression of TLR2 by HEK 293 SEAP cells was detected by immune-labelling using human anti-bovine CD282 antibody or mouse anti-human CD282 antibody conjugated with Alexa Fluor 647. HuCAL Fab-dHLX-MH antibody was used as a negative control.. A total of 10,000 events were recorded per construct by a FACS Calibur E3160 using Cell Quest Pro acquisition software (BD Biosciences, UK). Data was analysed using FlowJo V10 (FlowJo LLC, USA). Gating of SEAP cell population was based on forward-side scatter (A) and gating strategy was used for all constructs (B-G). Images represent the sum of three independent transient transfections per construct. The mean fluorescent intensity of cells from three independent transient transfections per construct is shown (H)

| | | |
|---|---|---|
| Number of scaffolds | 14729 | |
| Total size of scaffolds | 2658233619 | |
| Longest scaffold | 110669201 | |
| Shortest scaffold | 1000 | |
| Number of scaffolds > 1K nt | 14718 | (99.925%) |
| Number of scaffolds > 10K nt | 1747 | (11.861%) |
| Number of scaffolds > 100K nt | 345 | (2.342%) |
| Number of scaffolds > 1M nt | 193 | (1.310%) |
| Number of scaffolds > 10M nt | 69 | (0.468%) |
| Number of scaffolds > 47M nt | 10 | (0.068%) |
| Mean scaffold size | 180476 | |
| Median scaffold size | 2676 | |
| N50 scaffold length | 26027505 | |
| L50 scaffold count | 29 | |
| scaffold %A | 28.85 | |
| scaffold %C | 20.72 | |
| scaffold %G | 20.71 | |
| scaffold %T | 28.85 | |
| scaffold %N | 0.86 | |
| scaffold %non-ACGTN | 0.00 | |
| Number of scaffold non-ACGTN nt | 0 | |
| Percentage of assembly in scaffolded contigs | 97.4% | |
| Percentage of assembly in unscaffolded contigs | 2.6% | |
| Average number of contigs per scaffold | 1.7 | |
| Average length of break (>25 Ns) between contigs in scaffold | 2253 | |
| Number of contigs | 24799 | |
| Number of contigs in scaffolds | 11290 | |
| Number of contigs not in scaffolds | 13509 | |
| Total size of contigs | 2635523319 | |
| Longest contig | 3312623 | |
| Shortest contig | 48 | |
| Number of contigs > 1K nt | 23656 | (95.391%) |
| Number of contigs > 10K nt | 10149 | (40.925%) |
| Number of contigs > 100K nt | 5821 | (23.473%) |
| Number of contigs > 1M nt | 398 | (1.605%) |
| Number of contigs > 10M nt | 0 | (0.000%) |
| Number of contigs > 47M nt | 0 | (0.000%) |
| Mean contig size | 106275 | |
| Median contig size | 5493 | |
| N50 contig length | 522913 | |
| L50 contig count | 1496 | |
| contig %A | 29.10 | |
| contig %C | 20.90 | |
| contig %G | 20.89 | |
| contig %T | 29.10 | |
| contig %N | 0.00 | |
| contig %non-ACGTN | 0.00 | |
| Number of contig non-ACGTN nt | 0 | |

Supplementary Table 1: Scaffold Statistics

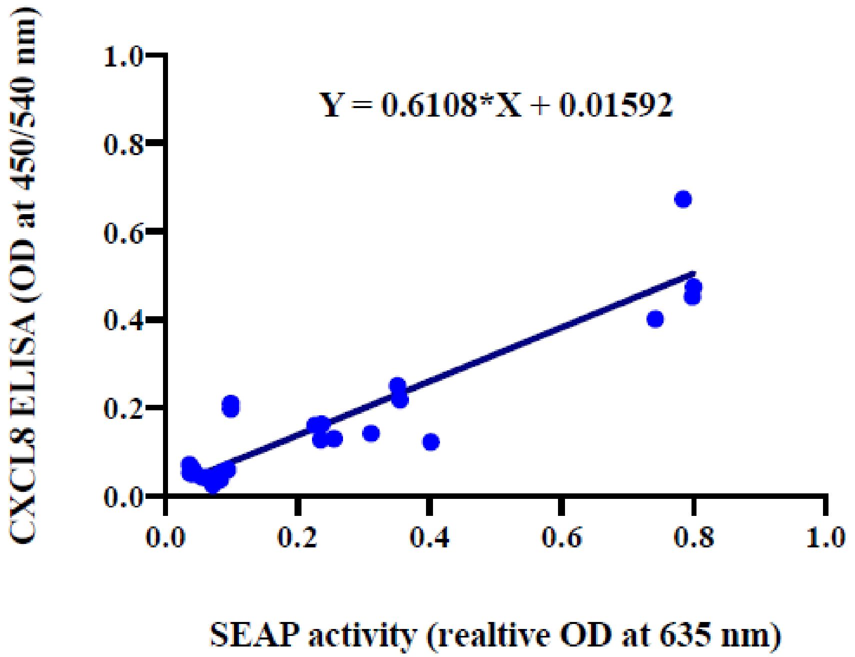


**Supplementary Figure S2: Correlation of ODs measured in SEAP reporter and CXCL8 ELISA assay**

ODs measured in the same supernatants of transfectants stimulated with FSL-1 at 100 ng/ml (Invivogen, USA), Pam$^3$CSK$^4$ at 1 mg/ml (Invivogen, USA) and their controls, measured in the SEAP assay and human CXCL8 ELISA assay using Quantikine Human CXCL8/IL-8 ELISA kit (R&D Systems, USA) were analysed for correlation between both assays. Spearman's correlation was implemented and found to be significant with a strong degree of correlation (r=0.7537, p<0.0001). To display this correlation, a dot plot fitted with Deming's regression line was drawn. Deming's regression line and equation are shown for one representative experiment. Data analysed and displayed with GraphPad Prism V8 (GraphPad Inc., USA)

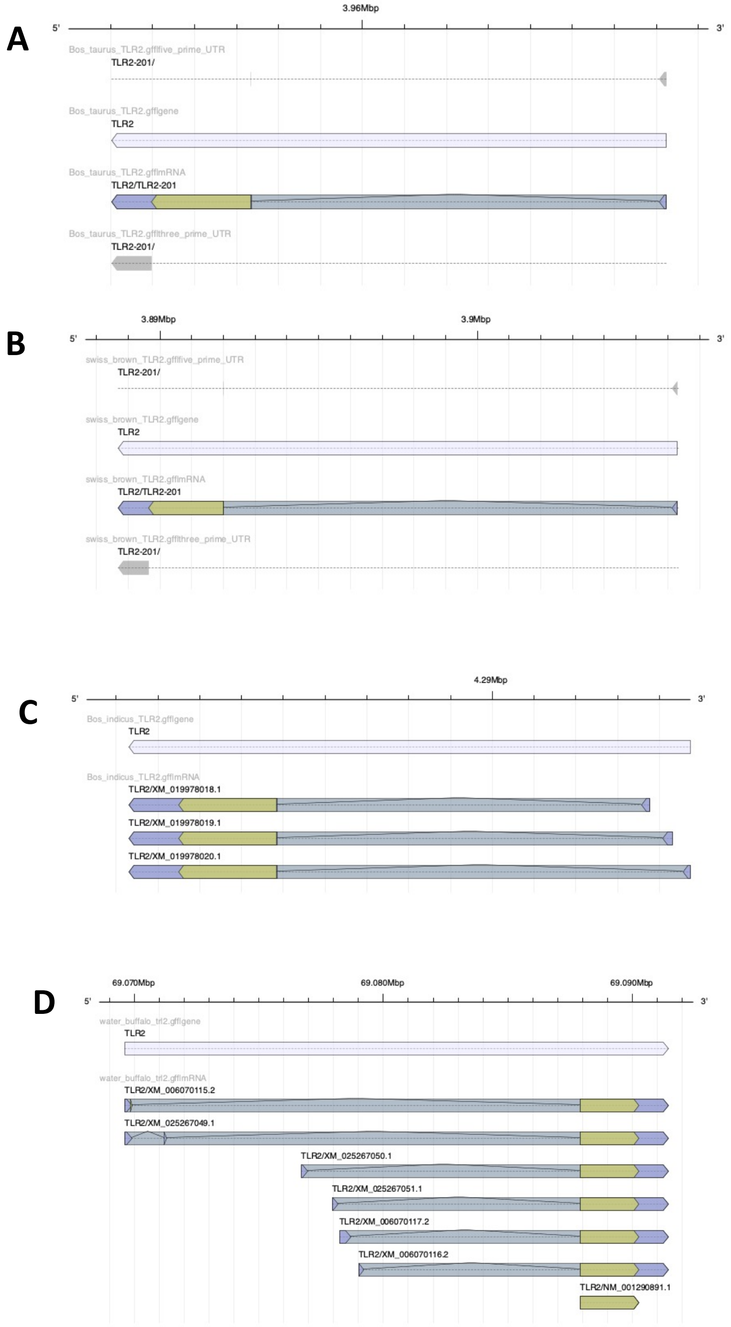


**Supplementary Figure S3: TLR2 Gene Models**

Gene models were created for TLR2 to represent *B. taurus* **(A)**, Brown Swiss **(B)**, *B. indicus* **(C)** and *B. bubalis* **(D)** identify the number of possible transcripts.

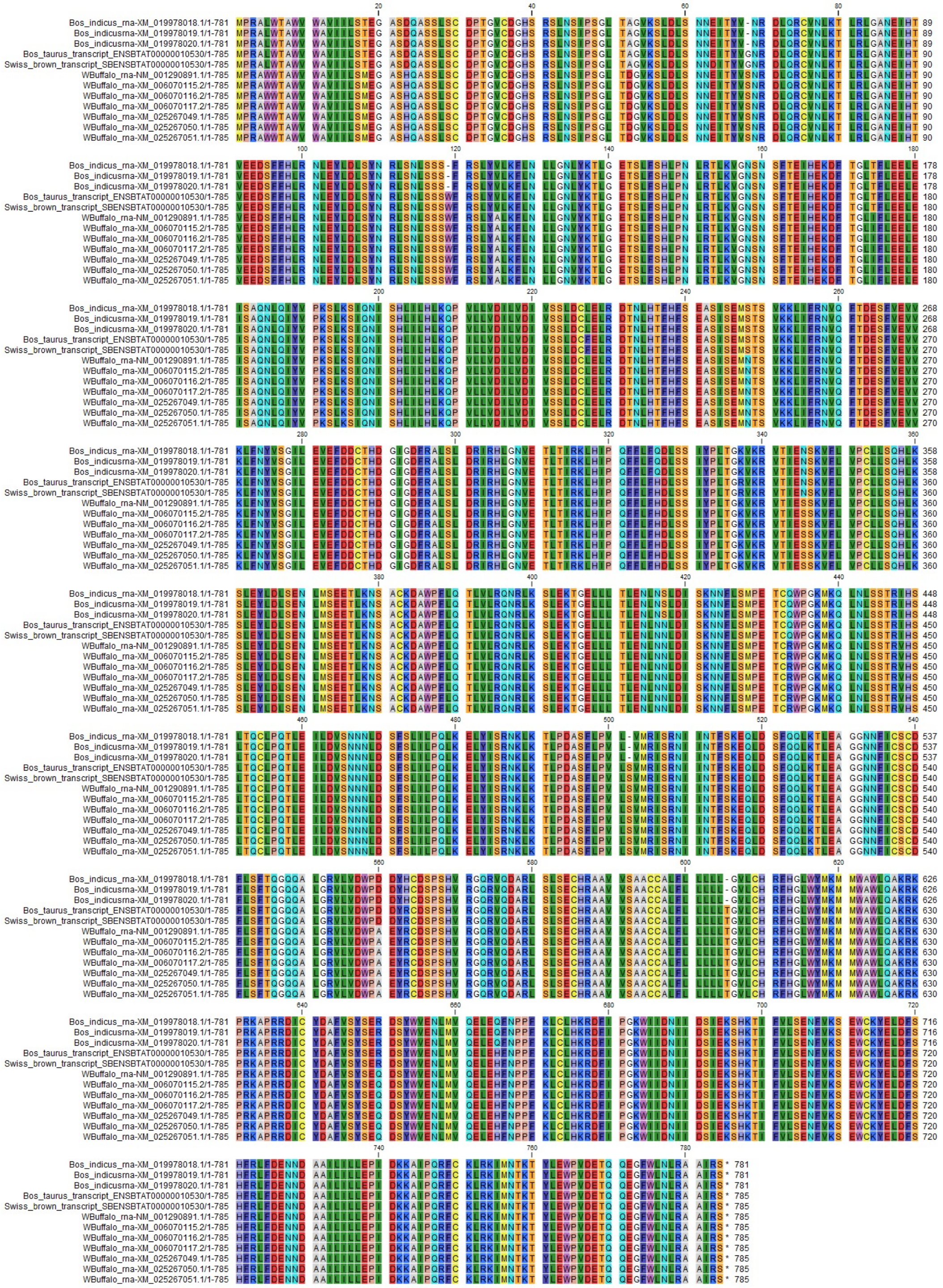

**Supplementary Figure S4: TLR2 CDS Alignment**

Coding sequences for TLR2 for each transcript defined in TLR2 gene models were extracted and aligned with ClustalW and translated to protein sequences.

**Notes:**

Sites where Brown Swiss differs from all others.
63 (D>E) GAG > GAT transversion
D=ASP Aspartic acid
E=GLU Glutamic acid

Sites where Brown Swiss and Bos taurus differ from all others

Sites where Brown Swiss and Bos taurus differ from Bos indicus Site (Bos taurus/Bos indicus)
211 (I>V)
227 (F>L)
337 (R>K)

Sites where Bos indicus differs from all others (B. indicus aa first)
119 (->W)
326 (Q>H)
417 (S>N)
502 (->S)
563 (H>R)
605 (->T)
665 (Q>H)

Variable sites
68 (-/G/S)
Bos indicus –
Bos taurus G
Brown Swiss G
Water Buffalo S
